# Computational Analysis of the Gut Microbiota-Mediated Drug Metabolism

**DOI:** 10.1101/2024.12.20.629788

**Authors:** Sammie Chum, Alberto Naveira Montalvo, Soha Hassoun

**Author notes:** Equal contributions (Sammie Chum, Alberto Naveira Montalvo). Availability and implementation: Code and dataset available at https://github.com/HassounLab/MDM.

## Abstract

The gut microbiota, an extensive ecosystem harboring trillions of bacteria, plays a pivotal role in human health and disease, influencing diverse conditions from obesity to cancer. Among the microbiota’s myriad functions, the capacity to metabolize drugs remains relatively unexplored despite its potential implications for drug efficacy and toxicity. Experimental methods are resource-intensive, prompting the need for innovative computational approaches. We present a computational analysis, termed MDM, aimed at predicting gut microbiota-mediated drug metabolism. This computational analysis incorporates data from diverse sources, e.g., UHGG, MagMD, MASI, KEGG, and RetroRules. An existing tool, PROXIMAL2, is used iteratively over all drug candidates from experimental databases queried against biotransformation rules from RetroRules to predict potential drug metabolites along with the enzyme commission number responsible for that biotransformation. These potential metabolites are then categorized into gut MDM metabolites by cross referencing UHGG. The analysis’ efficacy is validated by its coverage on each of the experimental databases in the gut microbial context, being able to recall up to 74% of experimental data and producing a list of potential metabolites, of which an average of about 65% are relevant to the gut microbial context. Moreover, explorations into ranking metabolites, iterative applications to account for multi-step metabolic pathways, and potential applications in experimental studies showcase its versatility and potential impact beyond raw predictions. Overall, this study presents a promising computational framework for further research and applications gut MDM, drug development and human health.

## 1. Introduction

The human gastrointestinal (GI) tract is colonized by ∼10^14^ bacteria belonging to at least several hundred species that are collectively termed the intestinal or gut microbiota [1]. This bacterial community is important for digesting dietary nutrients and extracting energy by fermenting carbohydrates indigestible by human enzymes [2]. The intestinal microbiota also impacts a wide range of functions in the GI tract [3] [4]. Beyond the GI tract, gut-brain [5], gut-lung, [6] and gut-liver [7] links have also been identified, thus highlighting the importance of the microbiota to human health and disease. Further, as microbiota-derived metabolites have been detected in circulation [8], alterations in the intestinal microbiota composition have been correlated to several diseases including obesity [9, 10], diabetes [11], cancer [12], and asthma [6].

A topic that is less explored, however, is characterizing the capacity of gut microbes to metabolize drugs, leading to drug activation, inactivation, and toxicity [13–15]. There are already documented studies on how gut microbes interfere with specific drugs. In some cases, such interference is by design, where a prodrug (non-active form) is transformed via microbial enzymes. An example is azo-bonded prodrugs of 5-aminosalicylic acid (5-ASA), including sulfasalazine, balsalazide, and olsalazine, that are used in the treatment of ulcerative colitis [16]. The azo bond links 5-ASA to a carrier molecule. When reaching the colon, azoreductases encoded by the gut microbiome cleave the azo bond to release 5- ASA. 5-ASA degradation rates are a function of gut composition and specificity on carrier molecules [17]. Other cases of interference are non-intentional and stomp researchers and clinicians for decades. One such example is cardiac drug digoxin [18], which was first isolated in the 1930s from the foxglove plant and became a drug in the 1960s. While the microbial species (*Eggerthella lenta*, formerly *Eubacterium lentum*) responsible for the degradation was identified in 1982 [19], the cardiac glycoside reductase genes encoding for this enzymatic degradation was only identified in 2013 and only in a few of *E. lenta’s* bacterial strains. These two examples and many others, e.g., ref [20–23], reflect the gut microbes’ enzymatic capacities for drug reduction, hydrolysis, acetylation/deacylation decarboxylation, dehydroxylation, demethylation, dehalogenation, and other reactions.

Concerns over microbiota-mediated drug metabolism (MDM) have prompted experimental testing the activity of drugs on panels of bacteria in isolation [24] and measuring isolate activities against a large panel of drugs [25]. Recently, the drug-metabolizing capacity of human fecal communities was challenged with 438 different compounds [26, 27]. A compounding challenge is that drug metabolization may occur through multiple steps that span microbes and the host [28], thus necessitating more complex experimentation. Complex experimental setups can now mimic the entire GI tract [29]. A further challenge is that microbial responses to drugs are a function of microbial community composition [30]. There are now databases that catalog drug-microbial interactions. The MDAD database provides a collection of microbe-drug associations [31], while the MASI (microbiota––active substance interactions) database offers a broad view of interactions between human microbiota and active substances such as drugs, dietary components, herbal products, probiotics, and environmental chemicals [32]. The MagMD (Metabolic action of gut Microbiota to Drugs) database was developed to specifically catalog interactions between gut microbiota and drugs, including detailed annotations on microbial enzymes, metabolites, and the effects on drugs [33]. MagMD data is collected from existing databases (e.g., PharmacoMicrobiomic [14], MASI [32]) and literature searches in PubMed using keywords related to gut microbiota and drug metabolism. To increase the coverage and annotation of MagMD, drug-microbe interactions were established by identifying microbes that had gene sequences with high similarity to sequences of known drug-metabolizing enzymes. For example, beta-glucosidase was responsible for 16,088 interactions between drugs and microbial species.

A complementary approach to experimental work and database curation is using computational approaches to analyze and predict MDM. Microbial promiscuity on drugs have been studied prior based on chemical similarity of drugs [34, 35], and by considering differences between substrate-product pairs of MetaCyc reactions [36]. A recent focus has been predicting microbe-drug associations. LCASPMDA combines a learnable graph convolutional attention network and iterative sampling ensemble strategy to infer latent microbe-drug associations [37], while GARFMDA combines graph attention networks and bilayer random forest to accomplish a similar task [38]. MDASAE uses stacked autoencoders to predict microbe-disease-drug associations [39]. These models, however, do not identify the enzymes responsible for the microbe activity on the drug, nor the metabolic product. There remains a need therefore to identify microbial metabolic products and the corresponding metabolizing enzymes, which in turn will guide experimentations and expedite MDM discoveries.

To identify drug-derived microbial metabolic products, this paper develops and evaluates a computational workflow, MDM, to predict putative products due to the activity of microbial enzymes on a query drug (**Figure 1**). To identify microbial enzymes, we crosslink KEGG Modules cataloged in the Unified Human Gastrointestinal Genome (UHGG) collection [40] to enzyme commission (EC) numbers within the Kyoto Encyclopedia of Genes and Genomes (KEGG) database [41, 42]. To predict MDM products, we utilize a rule-based method, PROXIMAL2 [43, 44] to predict such products. When specifying the enzyme’s Enzyme Commission (EC) number, PROXIMAL2 first identifies biotransformation rules relevant to the enzyme, and then applies them to the query drug to generate enzymatic products. The biotransformation rules are derived from reaction rules cataloged in the RetroRules database [45], which over 40,000 reaction rules extracted from public databases and encompass many drug-metabolizing reactions. We then use GNN-SOM [46], a site-of-metabolism predictor, to rank the resulting products. For evaluation, we curate several datasets from MagMD, MASI, and DrugBank [47], with and without known catalyzing enzymes, and assess the validity of using PROXIMAL2 to identify cataloged drug-derivative products and the corresponding microbial enzyme responsible for acting on the query drug. Further, we explore using all biotransformation rules in RetroRules to allow for the discovery of putative, yet currently undocumented, drug microbial metabolism. The contributions of the paper are:

- Developing the MDM computational workflow for predicting products of microbiota-mediated drug metabolism. Our workflow uses PROXIMAL2 integrated with biotransformation rules from RetroRules and microbial enzyme annotations from UHGG and KEGG, followed by ranking of predicted metabolites using GNN-SOM.
- Curating a list of known microbial enzymes by crosslinking KEGG Modules and the UHGG (see Supplementary Data 1).
- Curating two datasets to validate MDM predictions with gut-specific cases and two datasets with non-gut specific cases (see Supplementary Data 2). The datasets include 30 gut-specific, drug-product cases from MASI, MagMD, and the literature, and 465 non-gut specific cases of drugs from DrugBank metabolized by biological enzymes other than explicitly stated CYP enzymes.
- Demonstrating accuracy of MDM predictions using PROXIMAL2 with product recall rates of 73.91% on known gut-specific datasets, and 42.00% on non-gut specific datasets.
- Assigning microbial enzymes to 112 cases of documented drug-product relationships with unknown enzymes.
- Discovery that 29.86% of the non-gut specific dataset can be putatively metabolized by gut enzymes.

**Figure 1.**
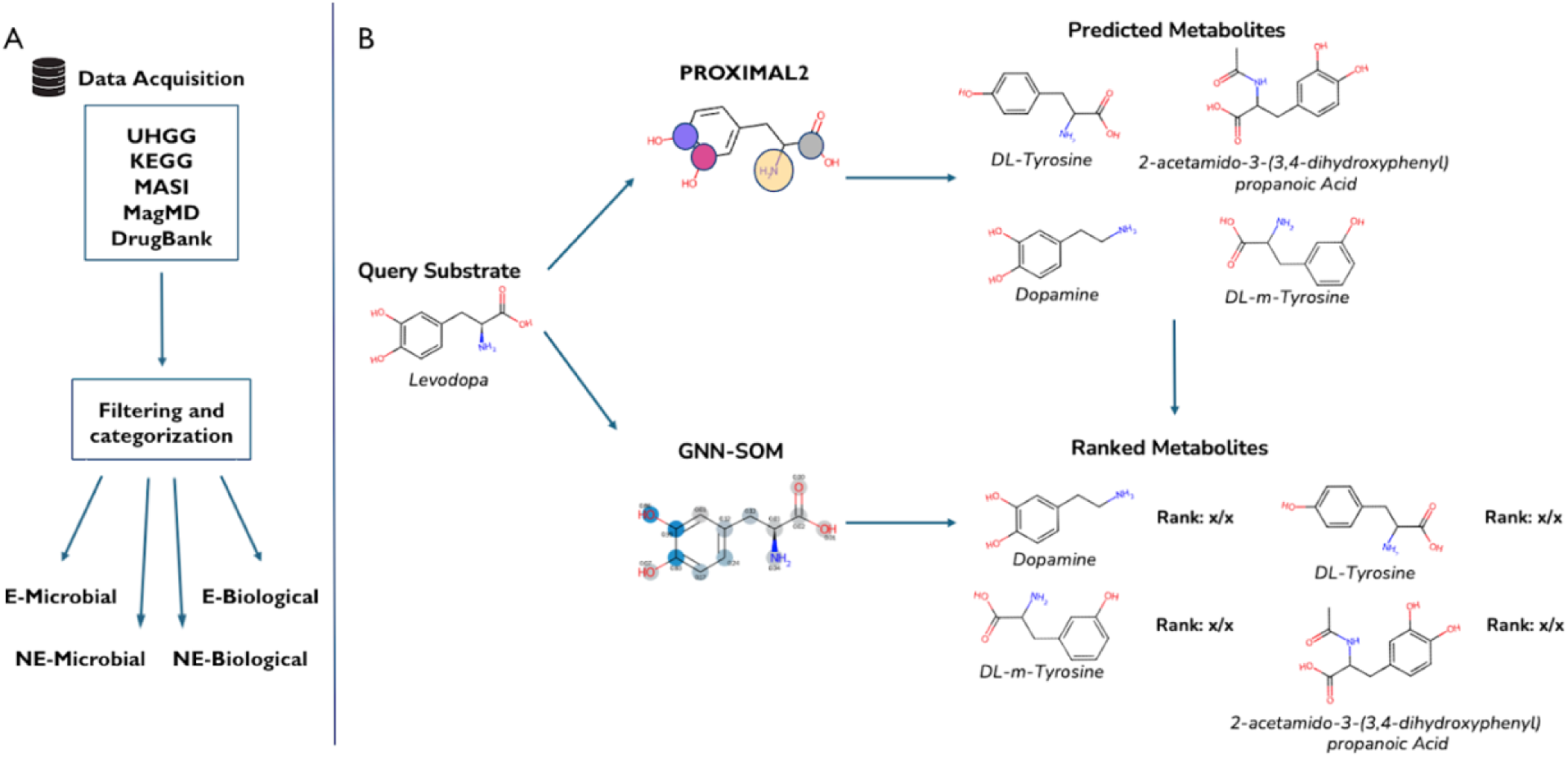
Overview of the computational workflow. (**A**) Data acquisition and curation by filtering reaction data and categorizing reactions into different datasets depending on the information available for that reaction. (**B**) MDM prediction via application of a metabolism prediction tool (PROXIMAL2) and ranking of predicted metabolites using GNN-SOM.

## 2. Methods

### 2.1 Data

Microbial enzymatic activity on drugs is cataloged in the MagMD and MASI databases. To explore if gut microbes can act on drugs utilizing enzymes whose function is similar to that of other biological enzymes, we curated additional datasets of drug-products from DrugBank. In total, we curated four datasets (**Table 1**). Datasets with known enzymes include an E prefix, whereas datasets with no known enzymes include an NE prefix.

**Table 1.** Four datasets were curated to validate and explore our methods, categorized by available information. “E” datasets contain known catalyzing enzymes, while “NE” datasets lack such information. “Microbial” sets contain MDM biotransformations that are known specific to the gut microbes, whereas “Biological” sets are not associated with the gut microbes. Each dataset includes counts of drugs, enzymes, and products.

|  | Interaction<br>known to<br>be gut-specific | Drug | Enzyme<br>known | Product | Number of<br>drugs | Number of<br>enzymes | Number of<br>products |
| --- | --- | --- | --- | --- | --- | --- | --- |
| E-Microbial | ☒ | ☒ | ☒ | ☒ | 20 | 16 | 23 |
| E-Biological | ☐ | ☒ | ☒ | ☒ | 100 | 100 | 100 |
| NE-Microbial | ☒ | ☒ | ☐ | ☒ | 7 | N/A | 7 |
| NE-Biological | ☐ | ☒ | ☐ | ☒ | 365 | N/A | 365 |

E-Microbial is a validation set for our method, consisting of drug-microbe-enzyme-product relationships that are known for microbial enzymes from MagMD and MASI and data from the literature that have not yet been added to these databases; E-Biological presents drug-enzyme-product relationships from DrugBank that are known for any biological enzyme. As data involving drug, microbe, metabolizing enzyme, and metabolic derivative products are limited, we expanded our curation efforts to include microbe-drug interactions with no known metabolizing enzyme. NE-Microbial and NE-Biological datasets consist of drug-product relationships from microbial and biological metabolism respectively, without a documented metabolizing enzyme.

The datasets were curated as follows. E-Microbial (**Table 2A**) was compiled from MagMD, MASI, and manual literature search. Each entry comprises a drug, microbe, enzyme name and corresponding EC number, and a metabolic derivative product. From MagMD and MASI, 16 unique cases of drug metabolism were extracted, with 5 overlapping cases between the two databases. Subsequently, manual literature search was utilized to identify cases that are not currently documented in the MagMD or the MASI databases. Seven additional cases of MDM were identified. E-Microbial therefore contains data for 20 unique drugs, with 23 unique drug-product pairs. The E-Microbial set is therefore used to validate the rule-based workflow’s ability in predicting metabolizing enzymes and their products.

**Table 2.**
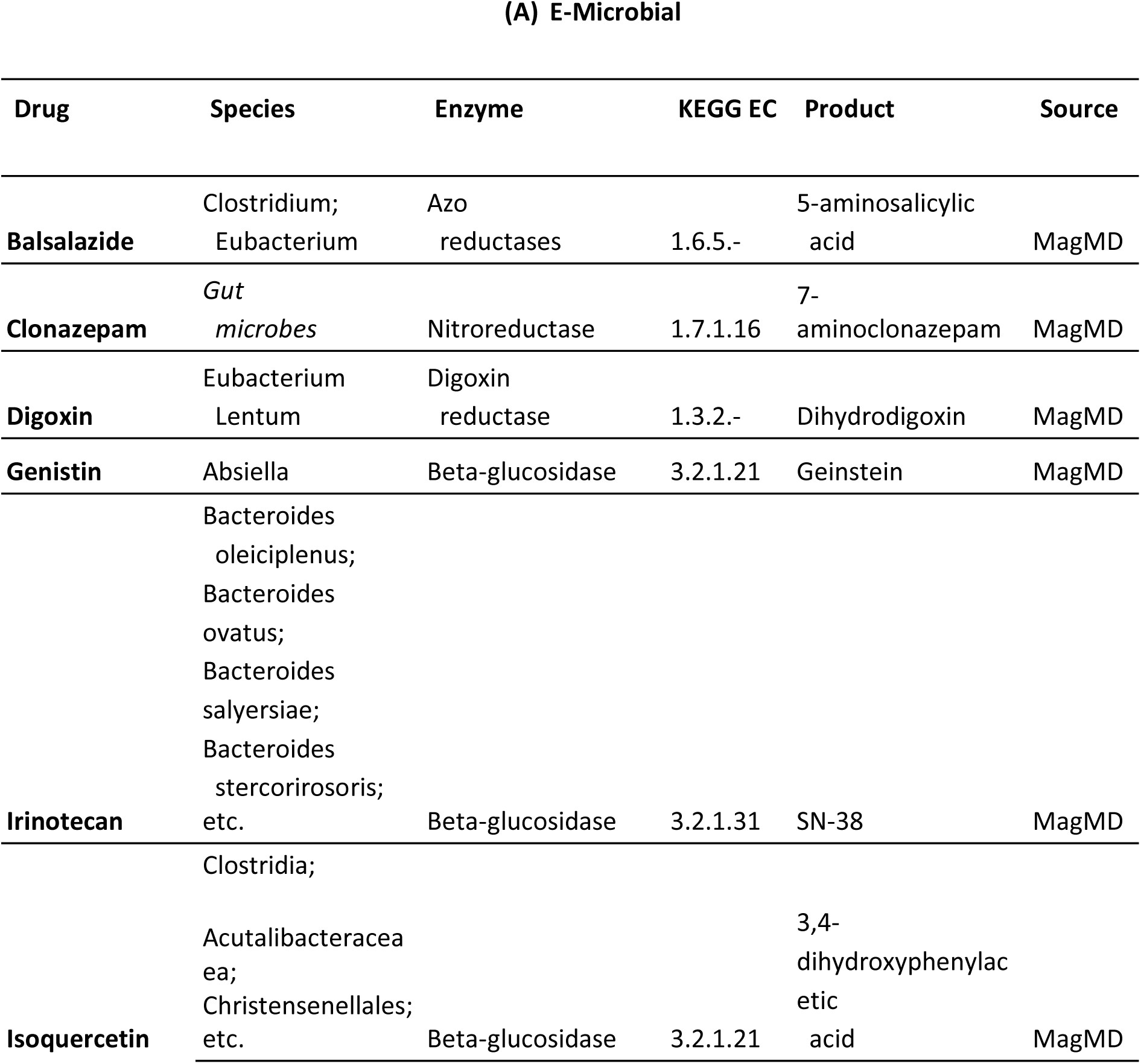
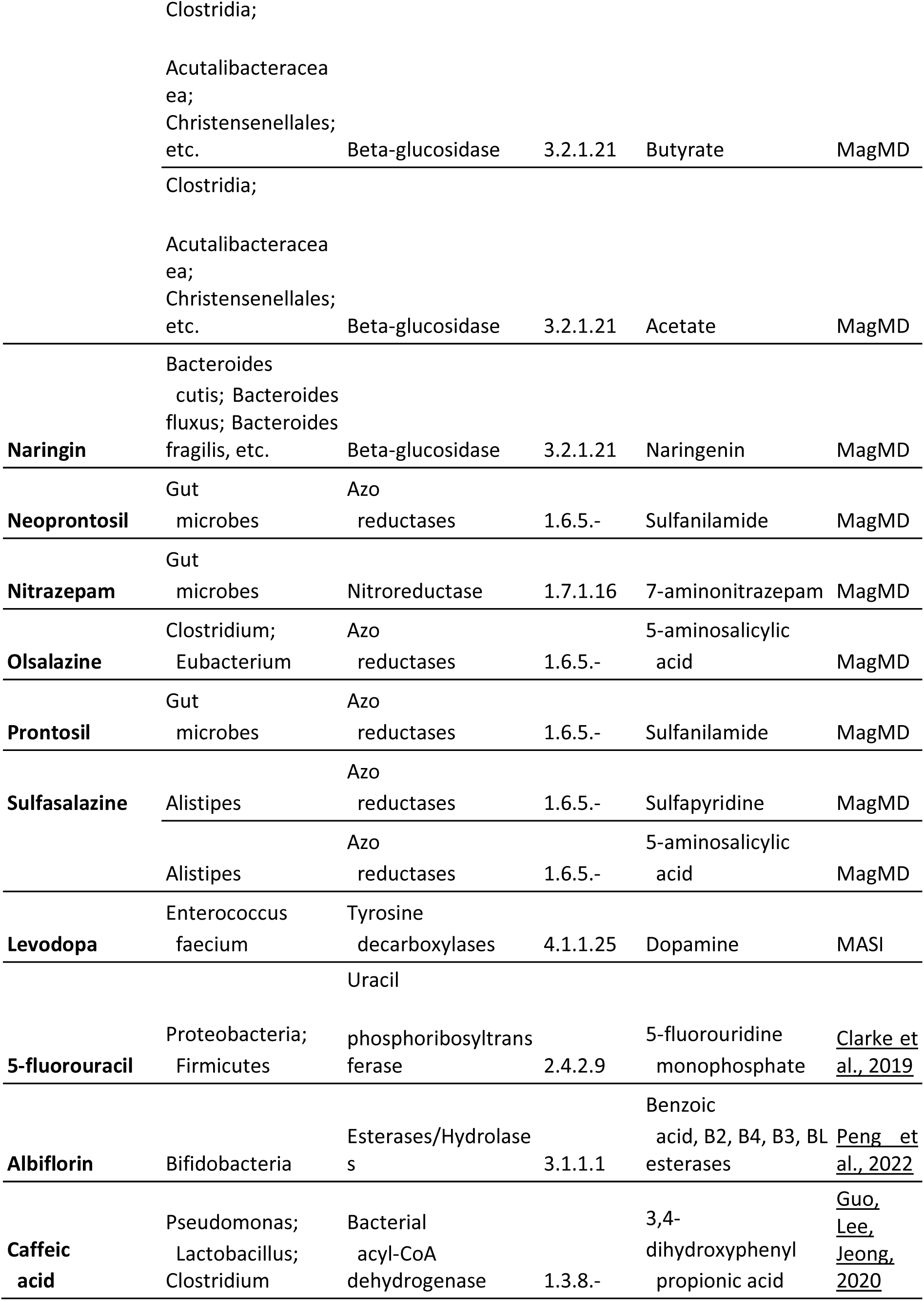
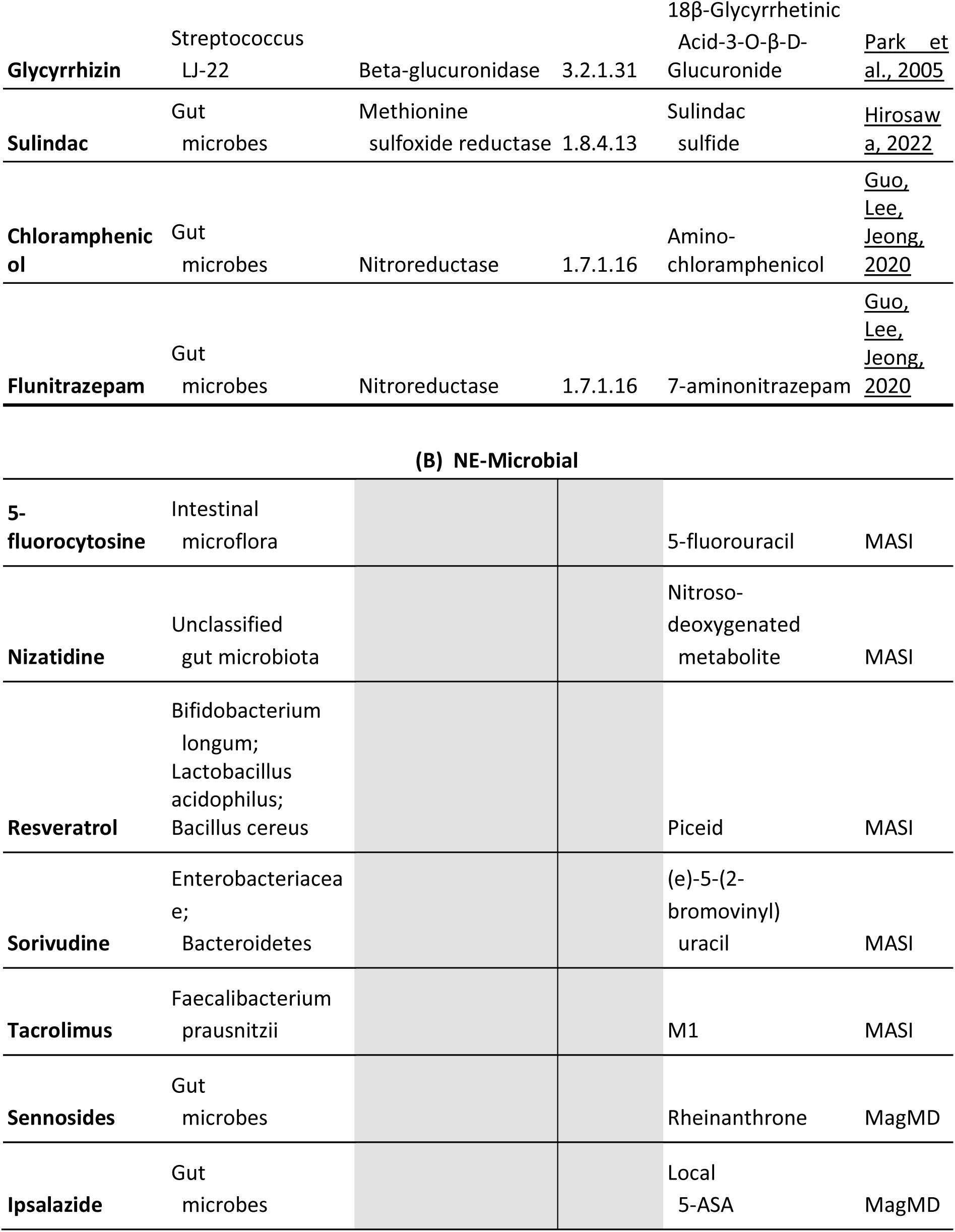
Gut-microbiome drug metabolism cases curated from MagMD, MASI, and manual search through the literature. **(A)** Cases with documented information on the drug, enzyme, and resulting metabolite. **(B)** Cases with documented information on the drug and resulting metabolite, but unknown acting enzyme or EC class. More detailed information is provided in Supplementary Table 1.

E-Biological was curated using DrugBank’s database of 3,703 drug metabolism reactions. Most participating enzymes of the DrugBank reactions are non-gut microbiome specific, with many of the enzymes to be known for liver metabolism. To remove some such reactions for a more balanced dataset, entries were filtered to exclude cases with keywords such as “liver”, “cytochrome”, and “CYP”. After filtering to exclude cases of non-gut microbiome-specific metabolism, empty entries, and entries without a documented enzyme, 100 reactions were designated as the E-Biological dataset. The E-Biological set is used to validate the rule-based workflow’s predictive ability of the appropriate metabolizing enzymes and products, and we also assert if the predicted enzymes are microbial.

The next two datasets, NE-Microbial and NE-Biological provide only drug-product pairs with an unknown metabolizing enzyme. NE-Microbial (**Table 2B**) was curated with cases specific to metabolism by the gut microbes, but unknown enzymes, from MASI and MagMD. Five cases were extracted from MASI and 2 from MagMD. Altogether, there are 7 drugs and 7 unique drug-metabolite pairs. Similarly, NE-Biological is composed of 365 DrugBank drug-product pairs with undocumented enzymes. In addition to validating if our method suggests the documented metabolic products, we report on gut microbiota enzymes that can metabolize these drugs.

### 2.2. PROXIMAL2 for predicting MDM

PROXIMAL2, a tool for the prediction of xenobiotic metabolism for CYP enzymes [43] and later to explore a wider range of enzymatic biotransformations [48], was repurposed for the prediction of gut MDM (**Figure 2**). Given a set of biotransformation rules from a database such as RetroRules [45], PROXIMAL2 creates lookup tables to catalog the biotransformation. Each entry in the table consists of a key and a value. The key consists of a putative reaction center (site-of-metabolism, SOM) and its adjacent neighboring atoms and distant adjacent atoms, i.e., a 2-hop neighborhood, all coded in KEGG atom types [41, 42]. The value describes the change to the reaction center and the neighbors, if applicable. Given a query drug and a specific enzyme, PROXIMAL2 will first identify the reaction rules associated with the enzyme. Then, PROXIMAL2 will iterate through the reaction rules and attempt to apply them to each atom within the query. PROXIMAL2 predicts several putative products, as shown in the example in **Figure 2**. As a standalone tool, PROXIMAL2 iterates over all reaction rules and attempts to apply them to each atom within the query. For the E-Microbial and E-Biological datasets where the catalyzing enzyme is known, the reaction rules associated with the known enzymes were used.

**Figure 2.**
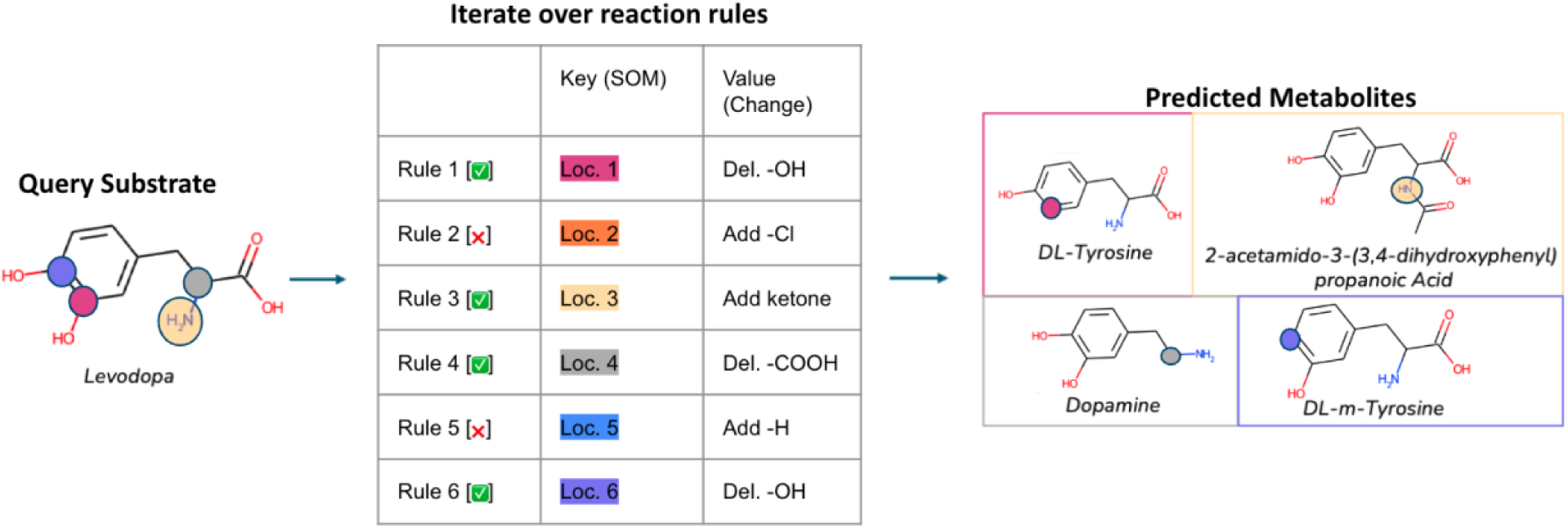
Schematic representation of PROXIMAL2 workflow. PROXIMAL2 iterates over atoms in the query substrate and applies any rule or change that has a corresponding key location that matches in the list of reaction rules. Rules 1, 3, 4 and 5 are matched with a location in the query whereas rules 2 and 5 were not matched. Locations are encoded using KEGG atom types. Each reaction rule is described by a key-value pair that encodes the site of metabolism (SOM) and the change that occurs, respectively.

### 2.3. Associating enzymes with microbial species

RetroRules biotransformations are associated with enzyme commission numbers (ECs) but are not linked to any species. We identify the subset of RetroRules biotransformations that can be attributed to microbial enzymes. We obtain a listing of known microbial enzymes as follows. The UHGG database [40] lists associations between microbes and their corresponding KEGG Modules, where each such module is manually curated to reflect a set of genes that operates as a functional unit. From the KEGG database, we extracted a set of enzymes (EC numbers) associated with these modules. We observed that of all the 42,307 biotransformations listed in RetroRules, 42.21% (17,858) are associated with enzymes relevant to the gut. As such, if an enzyme is contained within this set, then we can classify the corresponding biotransformation as due to the gut MDM. This list of microbial biotransformations, their relevant EC numbers, and corresponding species is available in **Supplementary Data 1 and 2.**

## 3. Results

### 3.1. Evaluating product and enzyme recall

We assess the capabilities of our approach in predicting microbial drug metabolism by evaluating its ability to recall known products, and enzymes, if known of microbial drug metabolism. Hence, PROXIMAL2 was applied to each drug present in the curated datasets using biotransformations associated with enzymes reported as responsible for such transformation (**Table 3A**). The resulting products are then compared against the products as reported in each of the corresponding datasets. Performance is measured using product recall, which is defined as the percentage of cases where the drug derivative product was reported by PROXIMAL2. We report the enzyme recall, which is the percentage of cases where the reported enzyme in the curated datasets led to the correct product. We also report on “both recall”, the percentage of cases where the correct product is produced due to the biotransformation rules associated with the documented enzyme. Indeed, there are products that PROXIMAL2 predicts are a result of the query drug interacting with the documented enzyme that do not match with the documented product in our curated dataset. An example is provided in **Figure 3**. Naringin, a flavonoid with anti-inflammatory properties, is correctly predicted by PROXIMAL2 to metabolize into naringenin when catalyzed by EC 3.2.1.21 (**Figure 3A**) through a bond cleavage. However, PROXIMAL2 also generates another metabolism prediction catalyzed by 3.2.1.21 with the introduction of a ring structure; a result of the same transformation applied in the opposite direction (**Figure 3B**).

**Figure 3.**
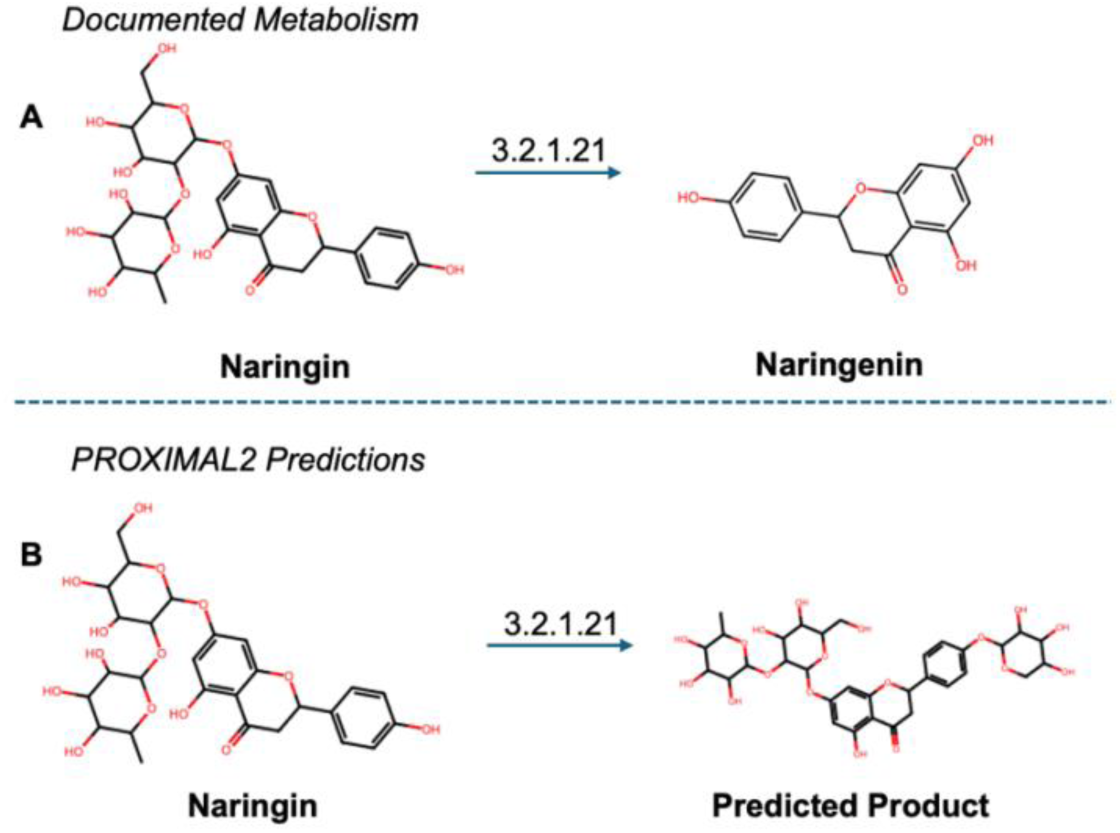
Two products derived by PROXIMAL of query drug naringin under EC. 3.2.1.21. (**A**) The documented transformation into naringenin (**B**) The predicted transformation to an undocumented metabolite.

**Table 3.** Results of applying PROXIMAL2 to predict microbial drug metabolism, reported using the recall metric on the product predicted by PROXIMAL2, on the enzyme (if specified for our curated dataset), and the recall of both product and enzyme, if specified. We also report the percentage of cases where the enzyme responsible for the biotransformation was associated with the gut microbes according to our UHGG analysis. **(A)** Using biotransformations that are *only* associated with the enzymes reported in our curated datasets. **(B)** Using all biotransformations available in RetroRules.

|  |  | (A) Biotransformation predictions due to known catalyzing enzymes |  |  |  | (B) Biotransformation predictions due to RetroRules |  |  |  |
| --- | --- | --- | --- | --- | --- | --- | --- | --- | --- |
|  |  | Product Recall (%) | Enzyme Recall (%) | Both Recall (%) | Percent in gut (%) | Product Recall (%) | Enzyme Recall (%) | Both Recall (%) | Percent in gut (%) |
| E-Microbial | any enzyme | N/A |  |  |  | 73.91 | 39.13 | 21.74 | 64.77 |
|  | enzymes associated with gut microbes | 47.83 | 39.13 | 21.74 | 100.00 | 65.22 | 39.13 | 21.74 | 100.00 |
| E-Biological | any enzyme | 36.00 | 35.00 | 14.00 | 56.41 | 42.00 | 35.00 | 14.00 | 60.26 |
|  | enzymes associated with gut microbes | 31.00 | 24.00 | 9.00 | 100.00 | 34.00 | 24.00 | 9.00 | 100.00 |
| NE-Microbial | any enzyme | N/A |  |  |  | 57.14 | N/A |  | 72.34 |
|  | enzymes associated with gut microbes |  |  |  |  | 42.86 |  |  | 100.00 |
| NE-Biological | any enzyme |  |  |  |  | 38.08 |  |  | 61.32 |
|  | enzymes associated with gut microbes |  |  |  |  | 29.86 |  |  | 100.00 |

When limiting PROXIMAL2 to utilize only the RetroRules entries linked to drug-metabolizing enzymes from on the E-Microbial dataset, we observe the following recall values: 47.83% of products were recalled, 39.13% of enzymes were recalled and 21.74% of cases with the documented enzyme predicting the documented product were observed (**Table 3A**)

When expanding the set of reference biotransformations on PROXIMAL2 to include the full RetroRules set, we see an increase in product recall to 73.91% for the E-Microbial set (**Table 3B**). However, only 64.77% of the predicted enzymes were relevant to gut MDM and so, when the products were filtered to only include these enzymes, the product recall decreased to 65.22%. Despite not being able to recall a large majority of products with just the enzymes listed in the dataset, when expanded to include all RetroRules, PROXIMAL2 is able to recall 65.22% of documented products in the E-Microbial dataset as a result of gut MDM. The increase in product recall when accounting for all biotransformation in RetroRules illustrates the ability of PROXIMAL2 to annotate products correctly beyond the biotransformations associated with the documented enzymes.

To assess the efficacy of PROXIMAL2 on non-gut MDM specific data, we evaluate the performance of PROXIMAL2 against the E-Biological dataset. On the run with biotransformation predictions due to enzymes documented in the E-Biological dataset, we observe a product recall of 36.00%, enzyme recall of 35.00% and both recall of 14.00% (**Table 3A**). However, pertaining to these predictions, only 56.41% of predicted enzymes were classified as relevant to gut MDM. As such, when we filter the results to only include gut MDM relevant enzymes the product, enzyme and both recalls decreased to 31.00%, 24.00% and 9.00%, respectively.

Expanding the set of reference biotransformations to the full extent of RetroRules, we see an increase in product recall for the E-Biological set to 42.00% (**Table 3B**). However, only 60.26% of the predicted enzymes were relevant to gut MDM and when the products were filtered to only include microbial enzymes, the product recall decreases to 34.00%. Similar to the E-Microbial dataset, we observe an increase in the product recall of the E-Biological dataset when accounting for all biotransformation in RetroRules. However, the recall metrics on this dataset were consistently lower than the corresponding metrics for the E-Microbial dataset. This result suggests that the biotransformations in the E-Biological dataset are not as clearly represented in the biotransformation rules within RetroRules.

As an additional evaluation for PROXIMAL2’s predictions, we assess the efficacy of PROXIMAL2 on the NE- Microbial and NE-Biological datasets which lack enzymatic information pertaining to each of the documented biotransformations. Specifically, we observe that PROXIMAL2 can recall 57.14% and 38.08% of products in the NE-Microbial and NE-Biological dataset respectively when run with the set of reference biotransformations in the full extent of RetroRules (**Table 3B**). Of these predicted biotransformations, 72.34% of enzymes for the NE-Microbial dataset and 61.32% of enzymes for the NE-Biological dataset were classified as relevant to gut MDM. When only considering these gut MDM specific enzymes, the observed product recall decreased to 42.86% and 29.86%. Consequently, we see that through these two examples, PROXIMAL2 can provide gut MDM specific enzyme recommendations for up to about 45% of biotransformations with unknown enzymes.

The ability to generate MDM predictions can also be leveraged to identify novel compounds measured in ex vivo studies, as was demonstrated prior for the CHO cell [49]. The application of PROXIMAL2 allows for the generation of a library of potential metabolites for experimentalists to compare chemically derived structures. Additional applications of comparing the similarity across derived structures with predicted structures provides confirmation, insight, and a potential route of metabolism with the generated EC prediction. There were two such structures in our datasets that we attempted to identify using PROXIMAL2. The derivative structures of the drug tacrolimus were experimentally determined utilizing MS and NMR, with the primary metabolite named M1, as the major metabolite, by researchers [33]. When applying PROXIMAL2, a library of 32 potential metabolites was generated, but none were identical to the experimental M1 structure. We also attempted to annotate the derivative product of the drug nizatidine. This derivative was classified as a nitroso-deoxygenated metabolite using HPLC/MS [14]. The application of PROXIMAL2 generated 3 potential metabolites, with none matching the experimentally derived structure.

We compare our PROXIMAL2 results with those obtained using BioTransformer, a computational tool for small molecule metabolism prediction [50, 51]. Instead of providing a set of reaction rules as input as in PROXIMAL2, BioTransformer users select predefined sets of enzymes to use for metabolism prediction. We used two such predefined sets. The “EC-based” set represents promiscuous metabolism using BioTransformer reaction rules, and hence comparable to PROXIMAL’s predictions using any enzymes provided via RetroRules. The “Human gut” set is used to predict xenobiotic metabolism by gut microbial enzymes, and hence comparable to PROXIMAL’s predictions using enzymes derived from RetroRules that are known to be associated with the gut microbes The results presented in **Table 4** for BioTransformer therefore aligns with the PROXIMAL2 results in **Table 3B**.

**Table 4.** Results of applying BioTransformer to predict products of microbial drug metabolism for the various datasets. The EC-based option in BioTransformer corresponds to utilizing all BioTransformer reaction rules, while the human gut option corresponds to utilizing enzymes associated by BioTransformer to the human gut. The N/A results are due to lack of specific enzyme information in NE datasets.

|  |  | Product Recall (%) | Enzyme Recall (%) | Both Recall (%) | Percent in gut (%) |
| --- | --- | --- | --- | --- | --- |
| **E-Microbial** | **EC-based** | 26.09 | 0.00 | 0.00 | 72.22 |
|  | **Human gut** | 52.17 | 39.13 | 17.39 | 63.16 |
| **E-Biological** | **EC-based** | 20.00 | 17.00 | 4.00 | 73.08 |
|  | **Human gut** | 17.00 | 16.00 | 4.00 | 61.54 |
| **NE-Microbial** | **EC-based** | 28.47 | N/A |  | 55.56 |
|  | **Human gut** | 14.29 |  |  | 60.00 |
| **NE-Biological** | **EC-based** | 15.89 |  |  | 76.47 |
|  | **Human gut** | 15.34 |  |  | 64.00 |

Recall values produced by BioTransformer are consistently lower than those produced by PROXIMAL2. Further, according to UHGG, enzymes predicted by BioTransformer to metabolize drugs in the human gut are relevant to gut MDM in 60- 64% of the analyzes cases. Reactions predicted by BioTransformer are derived for a set of 285 reference enzymes for EC-Based predictions and 53 reference enzymes for human gut microbial predictions [50]. Conversely, PROXIMAL2’s uses a set of 5,380 reference enzymes when utilizing all RetroRules, of which 2,500 are used as reference enzymes when analyzing gut microbes. Overall, PROXIMAL2 provides higher recall rates and utilizes a larger set of enzymes when predicting the gut MDM.

### 3.3. Case study 1: Iterative application of PROXIMAL2 to uncover the microbial metabolism for sennosides

Drugs are often metabolized through multiple enzymatic steps. For example, the drug levosimendan has been documented to first metabolize into an amino phenol pyridazinone metabolite, and then subsequently to an N-acetylated conjugate [52]. Here, we explore how the iterative application of PROXIMAL2 can elucidate multi-step drug metabolism.

The drug sennosides, used for treating constipation, is documented in the MagMD database to metabolize to the product rheinanthrone, as listed in NE-Microbial (**Table 2B**), but was not successfully predicted by PROXIMAL2. However, as shown in **Figure 4** and as documented prior [53], sennosides is metabolized in a series of biotransformations that first metabolizes to sennidin A, then to rheinanthrone (the active metabolite) and lastly rhein. We examine the iterative application of PROXIMAL2 to uncover this metabolizing pathway by considering each metabolite in this pathway as a query substrate to predict the correct subsequent product.

**Figure 4.**
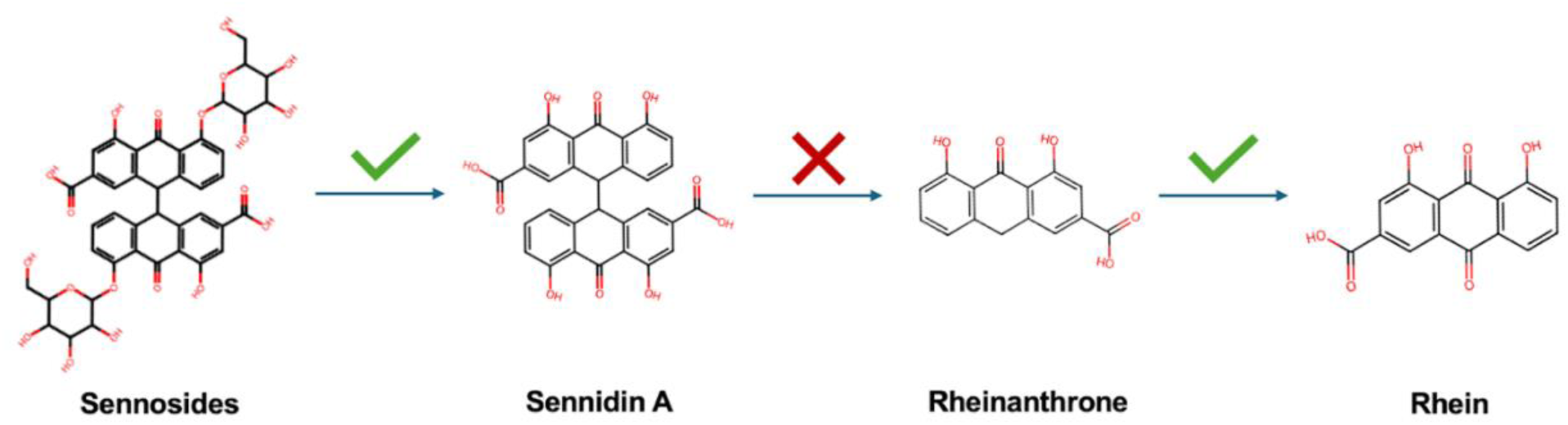
The biotransformation steps that the drug sennosides undergo to transform into product rheinanthrone, as documented in MagMD, and then to rhein. PROXIMAL2 was successful in predicting the metabolism of sennosides to sennidin A and rheinanthrone to rhein, but unable to predict the transformation from sennidin A to rheinanthrone.

The first product of this pathway, sennidin A, was correctly predicted by PROXIMAL2 as a product of sennosides through a hydrolysis reaction catalyzed by beta-glucosidase. Subsequently, sennidin A was expected to metabolize into rheinanthrone through a reductive carbon-carbon cleavage more commonly performed by Cytochrome P450 enzymes found in the liver. This product was not generated with PROXIMAL2 due to the lack of a similar transformation rule in the RetroRules database. Although carbon-carbon cleavages exist in RetroRules, none reflect similar molecular neighborhoods and therefore were not applied to the substrate. Lastly, when rheinanthrone was utilized as a query substrate, PROXIMAl2 was successful in generating the product rhein. In addition to product prediction, PROXIMAL2 was able to identify 3 corresponding enzymes likely to catalyze rheinanthrone: 1.13.12.20, 1.13.12.21, and 1.13.12.22. These enzymes share the first three digits of their EC number, 1.13.12.x, representing a class of oxidoreductases that incorporate one oxygen atom. This is illustrated by rheinanthrone’s transformation to rhein (**Figure 4**) with the introduction of a double bonded oxygen atom, further supporting the ability to predict an appropriate catalyzing enzyme.

### 3.4. Case study 2: Ranking predicted products of microbial metabolism on tamoxifen and 5- fluorocytosine

The application of biotransformation rules to a query drug results in many putative products. As some enzymes are more promiscuous than others, the number of products varies. For example, as shown in **Figure 5**, enzyme 1.14.14.1 acting on estradiol resulted in 8 products when applying either the subset of rules or all RetroRules. The variation in predicted products is a result of 6 explored sites of metabolism in estradiol by PROXIMAL2, in addition to different combinations of the reactive sites. The catalyzing enzyme, 1.14.14.1, is a form of monooxygenase within the class of oxidoreductases known for the incorporation or reduction of an oxygen atom. The large range of reactions this enzyme is known to catalyze, including hydroxylation, deamination, dealkylation, oxidation, and epoxidation, further contributes to the many putative products of estradiol metabolism. Excluding this case, there was an average of 2.36 products predicted per correct documented enzyme for E-Biological and 9.0 products for E-Microbial, emphasizing the need to discern the correct product even with knowledge of the catalyzing enzyme. When using all biotransformations within RetroRules, the average number of products is 7.0 for E-Microbial and 2.5 for E-Biological.

**Figure 5.**
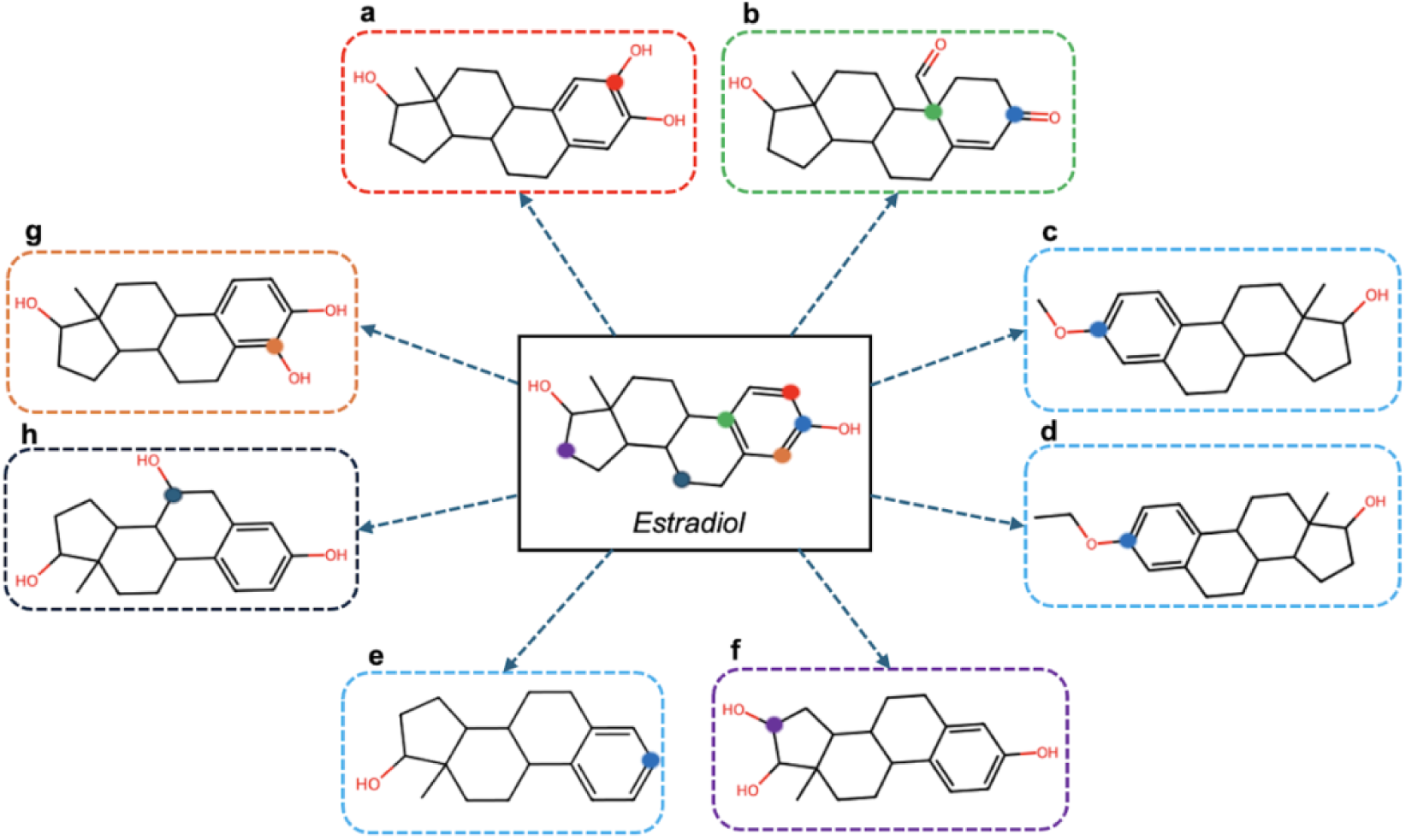
The eight products generated by the application of PROXIMAL2 to the query drug estradiol. The boxes around the molecules and the reaction centers are highlighted with the matching color to the query drug to indicate the site of metabolism, with an exception to product b containing two sites of metabolism.

Identifying the most likely product can be achieved via the application of GNN-SOM [46], a graph-based deep learning technique that can be leveraged to determine the most likely metabolites to be found in vivo. GNN-SOM is a graph neural net trained on all reactions from the KEGG RPAIR database by using RDM encodings, signifying the reaction center, difference, and match in each reactant pair. The final GNN-SOM consists of ten trained networks using the Chebyshev convolutional operator, with 2 or 3 layers, and a layer size of 512. GNN-SOM accepts a molecular fingerprint and EC number as input and returns a likelihood score at each atom of the compound, reflecting its likelihood to serve as a potential SOM.

By applying GNN-SOM to the substrate, we predict the likelihood of an atom within the substrate being the SOM (reaction center). If a product is derived based on a modification at a particular atom, this SOM likelihood is assigned as the likelihood of a product. We follow this product scoring method when an enzyme is specified, and all the derived products are realized via biotransformations associated with the specified enzyme. As the scores are normalized between 0 and 1, we also use this method to score the likelihood of the products when more than one enzyme generates the products, e.g., in the cases when the catalyzing enzyme is unknown. Two products may have the same score if the transformation occurs at the same SOM.

The application of GNN-SOM to one query drug use case, tamoxifen from E-Biological, was explored **(Figure 6).** Tamoxifen is used to treat breast cancer by blocking estrogen activity to mitigate tumor growth [54]. From PROXIMAL2, 5 products with 5 unique reaction centers were generated with the documented EC of 1.14.14.1. The documented metabolite was ranked first with a likelihood score of 0.58, and the second highest ranked metabolite scored 0.08. Reaction centers with higher likelihood scores than the known center, such as the nitrogen atom with a score of 0.78, were also labeled by GNN-SOM but no products were derived by PROXIMAL2.

**Figure 6.**
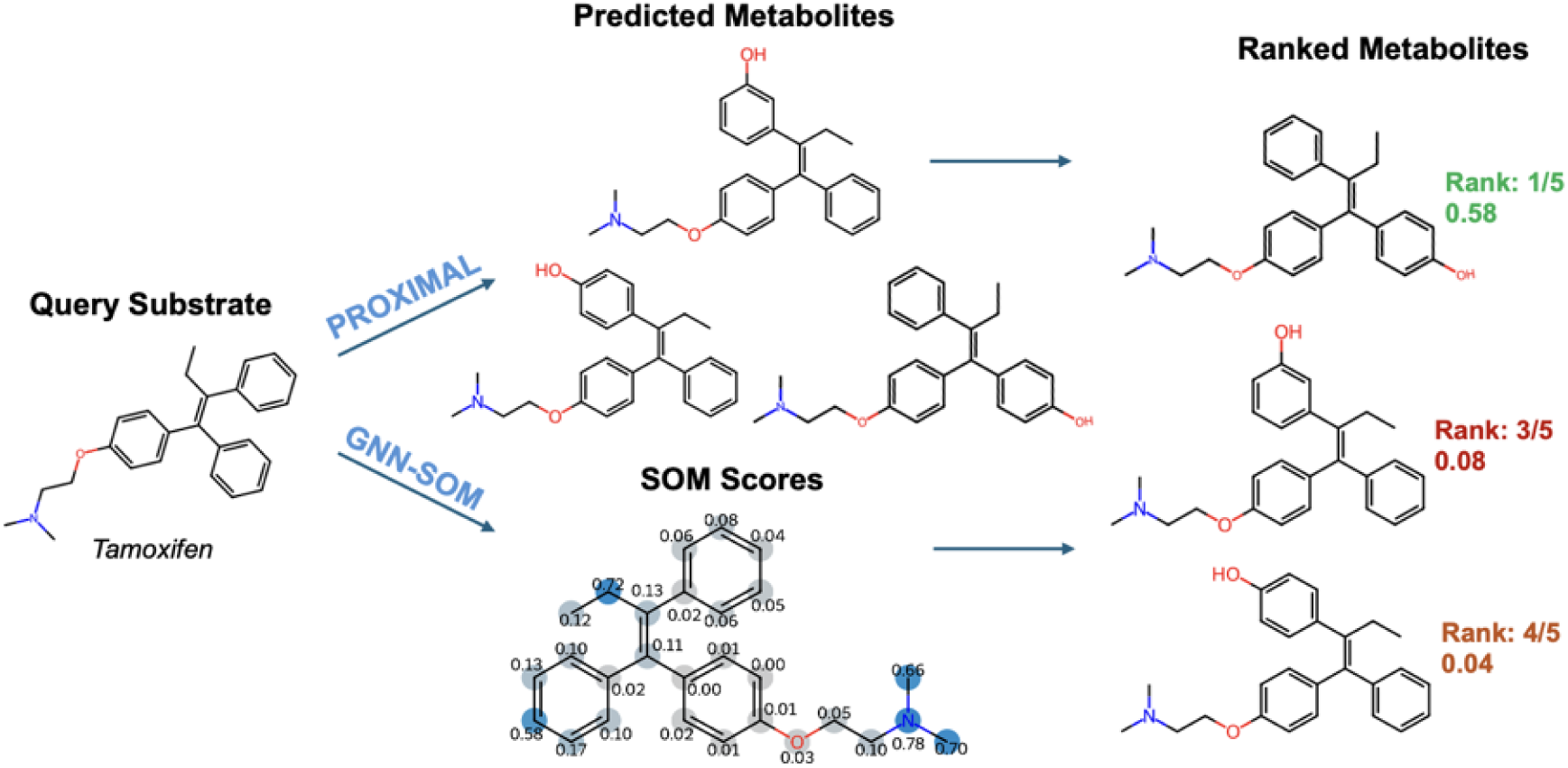
The analysis of tamoxifen from E-Biological with PROXIMAL2 and GNN-SOM to rank all predicted candidate metabolites. Three of the predicted metabolites are showcased, with the correct product ranked first out of five products with the documented EC.

The application of GNN-SOM-based ranking to another query drug, 5-fluorocytosine from NE-Microbial was also explored **(Figure 7)**. This drug is used as treatment for fungal infections [55]. PROXIMAL2 generated 12 unique products with 6 unique enzymes and 3 potential reaction centers. Upon analysis of all likelihood scores generated by drug and EC pairs, the top ranked product corresponded with the documented product of 5-fluorouracil with a likelihood score of 0.96 and was the only predicted product with the correct reaction center. The second and third ranked products exhibited significantly lower likelihood scores of 0.104 and 0.009, reflecting unlikely SOMs and GNN-SOM’s potential in determining the most likely products without a known EC. Although the current model of GNN-SOM is not trained on drug-specific candidates, the model demonstrates promising results if repurposed as a tool for processing PROXIMAL2 predictions.

**Figure 7.**
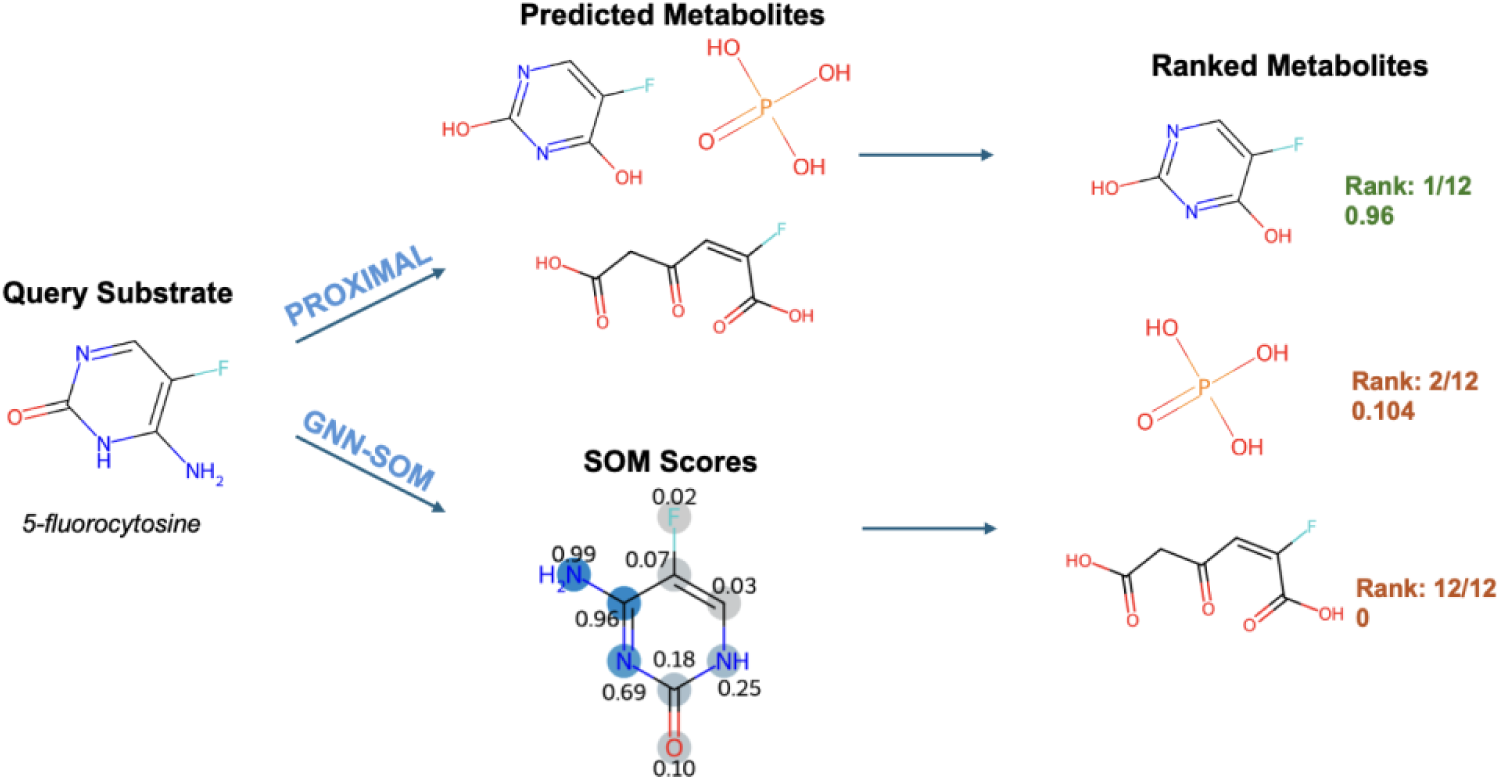
The analysis of 5-fluorocytosine from NE-Microbial with PROXIMAL2 and GNN-SOM to rank all predicted candidate metabolites. Three of the predicted metabolites are showcased, with the correct product, 5-fluorouracil, ranked first with a score of 0.96.

## 4. Discussion & Conclusion

This study explored gut MDM using computational methodologies, leveraging the PROXIMAL2 tool and integrating data from sources like the UHGG and KEGG. We developed a framework for predicting putative products of microbiota-drug-metabolism, validated against curated datasets, and analyzed for product/enzyme recall and enzyme prediction. Our findings elucidate PROXIMAL2’s efficacy in predicting both products and relevant enzymes in the gut microbial context. We accomplished the development of a robust prediction methodology, assessing PROXIMAL2’s performance across diverse datasets. Our analyses reveal insights into the complex interactions of gut MDM. Furthermore, by exploring iterative applications of PROXIMAL2 and envisioning potential applications in ex vivo experimental studies, we demonstrate the versatility and potential impact of our MDM pipeline in advancing drug discovery. These findings contribute to understanding gut MDM and pave the way for innovative approaches in deciphering this complex ecosystem. This has significant implications for human health and drug development, offering new avenues for research and application in the field.

A key consideration in evaluating the applicability of the MDM workflow is the balance between recall and the number of predictions generated. While precision alone does not fully capture predictive reliability, recall provides insight into how well-known metabolic transformations are recovered. The observed recall of ∼30% in certain cases reflect the challenge of capturing all enzymatic transformations with available biotransformation rules. Importantly, many PROXIMAL2 predicted products will fall outside our curated datasets, making it difficult to determine their validity without experimental validation. However, our approach is valuable as a hypothesis-generation tool, guiding experimentalists toward likely metabolic transformations rather than serving as a definitive predictor of MDM products. Future work incorporating confidence scoring is needed to prioritize predictions and improve interpretability for new drug applications.

A limitation of this study is the lack of species-level resolution in linking drug-metabolizing enzymes to specific gut microbial species. While we cross-referenced enzyme commission (EC) numbers with microbial gene annotations in UHGG and KEGG, the presence of an enzyme-associated gene does not necessarily confirm metabolic activity at the species level. Drug metabolism may require specific microbial strains with functional enzyme expression, metabolic cofactors, or interactions within the microbial community. Additionally, environmental factors such as gut pH, substrate availability, and microbial competition may further influence metabolic outcomes. Future work should aim to refine species-specific predictions by integrating metatranscriptomic data or experimental validation to determine active metabolic pathways in individual species.

While databases such as MASI and MagMD curated known microbial activities on drugs, there remains a need to uncover novel interactions between microbes and drugs. We investigated how rule-based methods like PROXIMAL2 can perform when challenged with a query drug and a microbial enzyme, or when challenged with a known drug-product pair to recover the microbial enzyme responsible for such transformation. The product recall rate of 73.91% on E-Microbial when using all biotransformations within RetroRules indicate either that the enzymatic transformation to produce the product is not available, or that multiple enzymatic steps are needed to generate the product thus reflecting inconsistencies in recording products within the database sources. Further, when comparing the 73.91% recall rate with the low recall rate of 47.83% for E-Microbial when using the named enzyme indicates that there are potential inaccuracies in recording the named enzyme in the literature or that the biotransformation rules associated with that enzyme are not complete within RetroRules. In either case, PROXIMAL2 may have shortcomings in applying complex transformations to generate the products, calling for improvements in product prediction

## 5. Acknowledgements

Research reported in this paper was supported by the Office of the Director of the National Institutes of Health under awards number R03OD036490 and R35GM148219. The content is solely the responsibility of the authors and does not necessarily represent the official views of the National Institutes of Health.

## Notes

https://tufts.box.com/s/xf4afk6araujo4jz6pe6301iuf19do5y

### Competing Interest Statement

The authors have declared no competing interest.

### Summary of Updates

The manuscript has been revised to enhance clarity, methodological transparency, and validation of results. To address the lack of species-level resolution in linking drug-metabolizing enzymes to specific gut microbes, the Discussion section now acknowledges this limitation and suggests future integration of metatranscriptomic data for improved species-specific predictions. Table 1 has been revised to include numerical values for the number of drugs, enzymes, and products in each dataset to improve readability and provide clearer insights into dataset composition. A benchmarking study has been conducted to compare PROXIMAL2 against BioTransformer, a widely used metabolism prediction tool. New results, presented in Table 4, highlight PROXIMAL2's higher recall rates and broader enzyme coverage. The Results and Discussion sections have been updated to provide a comparative analysis of the strengths and limitations of each tool. The workflow has been explicitly defined, and a new figure has been added to illustrate its structure. The revised Introduction and Contributions sections clarify that the methodology and dataset will be made openly available, with the code and dataset now accessible on GitHub. Further clarifications have been made regarding dataset curation and filtering. The Data section now provides additional details on how datasets were refined, including the rationale behind final dataset sizes. The filtering code is also available in the GitHub repository. Terminology and figures have been refined to improve precision. The term "model" has been replaced with "rule-based workflow," Figure 1's color scheme has been adjusted for consistency, and Figure 2's caption has been updated to clarify the distinction between documented and predicted transformations. Additionally, ambiguous phrasing regarding PROXIMAL2's predictions has been revised for clarity. The explanation of GNN-SOM's application has been improved. The revised Results section now explicitly describes its role in ranking predicted metabolic products based on site-of-metabolism likelihood. Case studies on tamoxifen and 5-fluorocytosine have been included to demonstrate the approach. In response to concerns about predictive reliability, the Discussion now contextualizes the workflow as a hypothesis-generation tool rather than a definitive predictor, emphasizing the need for future validation and confidence scoring. Lastly, the manuscript has undergone thorough proofreading by a native English speaker to improve readability and ensure clarity in sentence structure and scientific communication.

